# *alms1* regulates the immune response and brain ageing in zebrafish

**DOI:** 10.1101/2025.04.19.639742

**Authors:** Brais Bea-Mascato, Luis Méndez-Martínez, Carolina Costas-Prado, Laura Guerrero-Peña, Paula Suarez-Bregua, Josep Rotllant, Diana Valverde

**Affiliations:** CINBIO, University of Vigo, 36310 Vigo, Spain; Rare Diseases and Paediatric Medicine Research Group, Instituto de Investigación Sanitaria Galicia Sur (IIS Galicia Sur), SERGAS-UVIGO, Vigo, Spain; Aquatic Biotechnology Lab, Institute of Marine Research, Spanish National Research Council (IIM-CSIC), Vigo, Spain

## Abstract

The *ALMS1* gene plays a crucial role in maintaining cellular homeostasis through its involvement in primary cilium assembly, cytoskeletal regulation, and signalling pathways such as NOTCH and TGF-β. Pathogenic variants in *ALMS1* are associated with Alström Syndrome (ALMS), a multi-systemic ciliopathy characterised by neurosensory deficits, metabolic disorders, and multi-organ fibrosis. To better understand the tissue-dependent role of *ALMS1*, we utilised CRISPR/Cas9 technology to develop a zebrafish model with *alms1* depletion. Multi-tissue transcriptomic profiling revealed that *alms1* depletion has pleiotropic effects on gene expression, with the brain and eyes displaying the most pronounced transcriptomic alterations, including disrupted ciliary function and immune dysregulation. Inflammatory and innate immune pathways along with glutamatergic synapse-related processes were significantly affected in the brain and eyes but with different gene expression signatures. The analysis further highlights tissue-specific processes, primarily associated with organ dysfunction. Additionally, our findings underscore the role of *alms1* in regulating age-associated gene expression profiles in the brain, suggesting a link between ciliary dysfunction and accelerated brain ageing. Comparative analyses with Bardet-Biedl Syndrome iPSC models revealed shared pathways, reinforcing the potential of ciliopathies as models for ageing-related disorders. This study provides novel insights into the tissue-specific functions of *alms1* and the molecular mechanisms underlying ALMS, paving the way for the development of targeted therapeutic strategies.

## INTRODUCTION

The *Alström syndrome 1 gene (ALMS1)* encodes a structural protein localized in the centrosomes [1]. This protein is responsible for maintaining cohesion between the two centrioles that form the primary cilium’s basal body. The primary cilium is a cellular organelle involved in signal transduction from the external environment to the cell interior [2–7]. Over time, the functions of the *ALMS1* gene have been associated with the regulation of the α-actin cytoskeleton, the assembly and morphology of the primary cilium, and the modulation of signalling pathways mediated by this organelle, including the NOTCH and TGF-β pathways [2,5,8–10]. In recent years, the primary cilium has emerged as a key player in the pathogenesis of various diseases, including cancer, polycystic kidney disease, and autoimmune disorders [11–19]. The dysfunction of one or more ciliary genes has been associated with a diverse group of syndromes, collectively known as ciliopathies, with 55 distinct disorders currently recognised [12,20].

Alström syndrome (ALMS, OMIM #203800) is a ciliopathy caused by pathogenic variants in the *ALMS1* gene [21]. Like most ciliopathies, this syndrome affects multiple organs. The most common phenotypic manifestations of this syndrome include progressive vision loss, which can ultimately lead to total blindness; hearing impairment; metabolic disorders, such as obesity or type 2 diabetes mellitus (T2DM); dilated cardiomyopathy; or multi-organ fibrosis, leading to hepatic, renal or pulmonary dysfunction [22–24]. The metabolic syndrome observed in these patients, along with its associated complications, such as obesity and type 2 diabetes mellitus (T2DM), represents one of the most extensively studied phenotypic manifestations in the scientific literature [25–30]. Nevertheless, the role of the *ALMS1* gene in most of the phenotypes observed in these patients remains unclear.

The multi-systemic nature of ALMS, combined with its low incidence of 1 in 1,000,000 and the consequent difficulty in obtaining samples, poses a significant challenge to its study [31,32]. In previous studies conducted in our laboratory, cellular models of retinal pigment epithelium (RPE), fibroblasts and tumour cell lines were used to identify processes potentially altered by *ALMS1* gene inhibition (siRNA and KO) and their relevance to the multi-organ fibrosis observed in these patients [8,9,33,34]. However, the resolution of this approach limits our ability to investigate processes at the cellular and tissue level. Accordingly, in this study, we developed a zebrafish animal model to investigate the tissue-specific implications of *ALMS1* gene depletion. Previous studies have documented the ability of this organism to replicate the ALMS phenotype [35]. Moreover, zebrafish has been widely used to investigate various ciliopathies, with a particular focus on ALMS [36–41]. The role of the *ALMS1* gene appears to be highly conserved across species, as evidenced by the ability of mouse models to replicate the metabolic phenotype observed in humans [42]. Thus, tissue-specific gene expression profiling in these models represents a promising approach to elucidating the context-dependent regulation of the *ALMS1* gene. Generating these enriched datasets will facilitate the prioritisation of the organ in which the *ALMS1* gene plays the most relevant role, thereby enabling a comprehensive understanding of the disease. This will contribute to the development of innovative and more effective therapeutic strategies for ALMS.

## MATERIALS AND METHODS

All experiments in this study were performed following all the principles outlined in the European Animal Directive (2010/63 UE) for the protection of experimental animals. Ethical approval was obtained from the Institutional Animal Care and Use Committee of the IIM-CSIC Institute and Xunta de Galicia in accordance with the National Advisory Committee for Laboratory Animal Research Guidelines licensed by the Spanish Authority (RD53/2013). The approval code is (ES360570202001 / 18 / FUN. 01 / BIOL AN. 08/JRM).

### Generation and analysis of *alms1* knockout mutants

The mutant line, carrying non-functional alleles for the *alms1* gene, was generated on a wild-type genotype background (TU). Using the CRISPR/CAS9 editing technique, two single-guide RNAs (gRNAs) were microinjected alongside the mRNA sequence for the CAS9 protein **(Supplementary Table 1)**. The gRNAs were identified and analysed using the Benchling platform **(Supplementary Table 2)**.

One-cell stage embryos were employed to generate the F0 generation, which consisted of genetic mosaics. The line was then crossed with wild-type individuals, resulting in the F1 progeny, of which a total of 47 fish were genotyped. From this generation, three non-functional alleles (NFA) were selected **(Supplementary Table 3)**. Subsequently, the heterozygous individuals carrying these NFAs were crossed with one another to produce homozygous individuals of the F2 generation, which were then cultivated for three months until they reached reproductive age.

Following the crossing and screening process, homozygous individuals for the three different alleles were obtained **(Supplementary Table 4).** In subsequent experiments, only homozygous F2 individuals carrying one of the three selected alleles were used, while F2 individuals with a wild-type genotype served as controls.

### FINCLIP genotyping

Genotyping of each generation was performed through DNA extraction from the caudal fin. The tissue sample was transferred in a 1.5mL tube containing 90µL of 50mM NaOH. The samples were initially incubated on a thermoblock for 30 minutes at 95°C. Subsequently, 10µL of 1M TRIS (pH 8) was added. The samples were stored at 4°C until polymerase chain reaction (PCR) amplification was conducted.

On the following day, a PCR was carried out to amplify exon 4 of the *alms1* gene, using the previously extracted genetic material **(Supplementary Table 1)**. The experiment was performed according to the protocol described by Bea-Mascato *et al.* [31]. Amplification products were validated on a 2% agarose gel with 0.05% ethidium bromide (EtBr) before being sequenced via Sanger sequencing.

The resulting sequences were subsequently analysed using the Inference of CRISPR Edits (ICE) analysis tool [43]. Three non-functional alleles (NFA) were selected in the F1 generation based on individuals exhibiting a knockout (KO) score of 50% (i.e., one null allele and one wild-type allele).

### Sample extraction and processing

Both *alms1* knockout individuals and their wild-type siblings were sacrificed at nine months of age. The specimens were euthanised with MS222, then dissected, and the tissues of interest (eyes, brain, and liver) were transferred to 1.5 mL tubes containing 750µL of TRIzol (Invitrogen, USA). To prevent degradation, the samples were maintained on ice throughout the process.

The tissue was initially disaggregated using scissors, treated with RNase away (Invitrogen, USA), and subsequently lysed by sonication with a Branson Sonifier SFX 250 (Branson Ultrasonics, USA). The sonication protocol consisted of 10 pulses of 5 seconds with 1-second rest between each pulse, at a wave amplitude of 20%. The protocol was repeated three times for each sample, with the samples kept on ice for three minutes following each sonication.

The samples were then centrifuged at 4°C and 12,000 rpm for 10 minutes. The supernatant obtained after centrifugation was transferred to Phase Lock Gel Heavy 2mL columns (Quantbio, USA), where 200µL of chloroform was added. The samples were centrifuged for 15 minutes at 12,000 rpm. The resulting supernatant was transferred to fresh 1.5 mL tubes, where it was mixed with an equal volume (1:1 v/v) of 70% ethanol. The mixture was then transferred to QIAGEN RNeasy columns (QIAGEN, Germany), and RNA extraction was performed according to the manufacturer’s protocol. The samples were eluted in 30 µL of RNase-free water (QIAGEN, Germany).

The RNA concentration was determined using the Nanodrop 2000c (Thermo Fisher Scientific, USA) and the Qubit 4 fluorometer (Invitrogen, USA). RNA integrity was assessed with the RNA IQ assay Kit (Invitrogen, USA) and all samples exhibited an IQ value greater than 7, indicating satisfactory integrity.

### Transcriptomic analysis of messenger RNA

For library preparation, the DNBSEQ eukaryotic stranded transcriptome library preparation protocol was followed according to the manufacturer’s instructions. The resulting libraries were sequenced on a DNBSEQ-G400 platform (BGI genomics, Hongkong), generating 100 base pairs (bp) paired-end reads.

A total of 18 samples were processed using the DNBSEQ platform, with an average data yield of 4.53GBs per sample. Initially, the Fastq files were filtered using SOAPnuke (v1.5.2; BGI) with the following parameters: *-l 15 -q 0.2 -n 0.05*. The filtered reads were then aligned to the *Danio rerio* genome (GCF_000002035.6_GRCz11, NCBI) using Bowtie2 (v2.2.5) with the parameters: *-q -- sensitive --dpad 0 --gbar 9999999999 --mp 1,1 --np 1 --score-min L,0,-0.1 -p 16 -k 200*.

Finally, gene expression levels were quantified using RSEM (v1.2.8) to generate count matrices, applying the parameters: *-p 8 --forward-prob 0 --paired-end*. The average gene set alignment ratio among the samples was 65.09%; a total of 25,702 genes were detected.

Differential expression analysis was performed using DESeq2, with an adjusted p-value (FDR) threshold of ≤ 0.05 and |log₂FC| ≥ 0.5.

Downstream analysis was conducted using over-representation analysis (ORA) with both Kyoto Encyclopaedia of Genes and Genomes (KEGG) and Gene Ontology (GO) term pathways. Subsequently, gene set enrichment analysis (GSEA) was performed using the same pathway databases to determine whether these processes were over- or under-activated.

For the analysis of GTEx enrichment and gene expression signatures associated with aging, Fisher’s exact test was applied. An adjusted p-value (FDR) threshold of <0.05 was used to identify statistically significant results.

## RESULTS

### Genotyping of non-functional alleles

To generate a zebrafish *alms1* knockout model, we first searched the zebrafish genome for orthologues of the human gene. A single zebrafish *alms1* gene was identified and verified by synteny analysis to detect any unannotated paralogues **(Figure S1)**.

We targeted exon 4 of *alms1* using the CRISPR/Cas9 gene editing system to introduce different pathogenic variants and characterise different non-functional alleles (NFA) **(Figure 1A, B)**. Previous studies confirmed that pathogenic variants in this exon can recapitulate the Alström syndrome (ALMS) phenotype in zebrafish [44]. Three heterozygous lineages with distinct NFAs were established **(Figure 1B, Figure S2A).** NFA1 carried a 5 bp deletion **(Figure S2A, B)**. NFA2 carried a 16bp deletion, while NFA3 carried a 7bp deletion **(Figure S2A, C, D)**. All these mutations resulted in a frameshift and a premature stop codon at amino acid positions 388, 383 and 386, respectively **(Figure S2E)**. Unfortunately, it was not possible to obtain homozygous individuals for NFA2, therefore the downstream analysis continued with NFA1 and NFA3.

**Figure 1.**
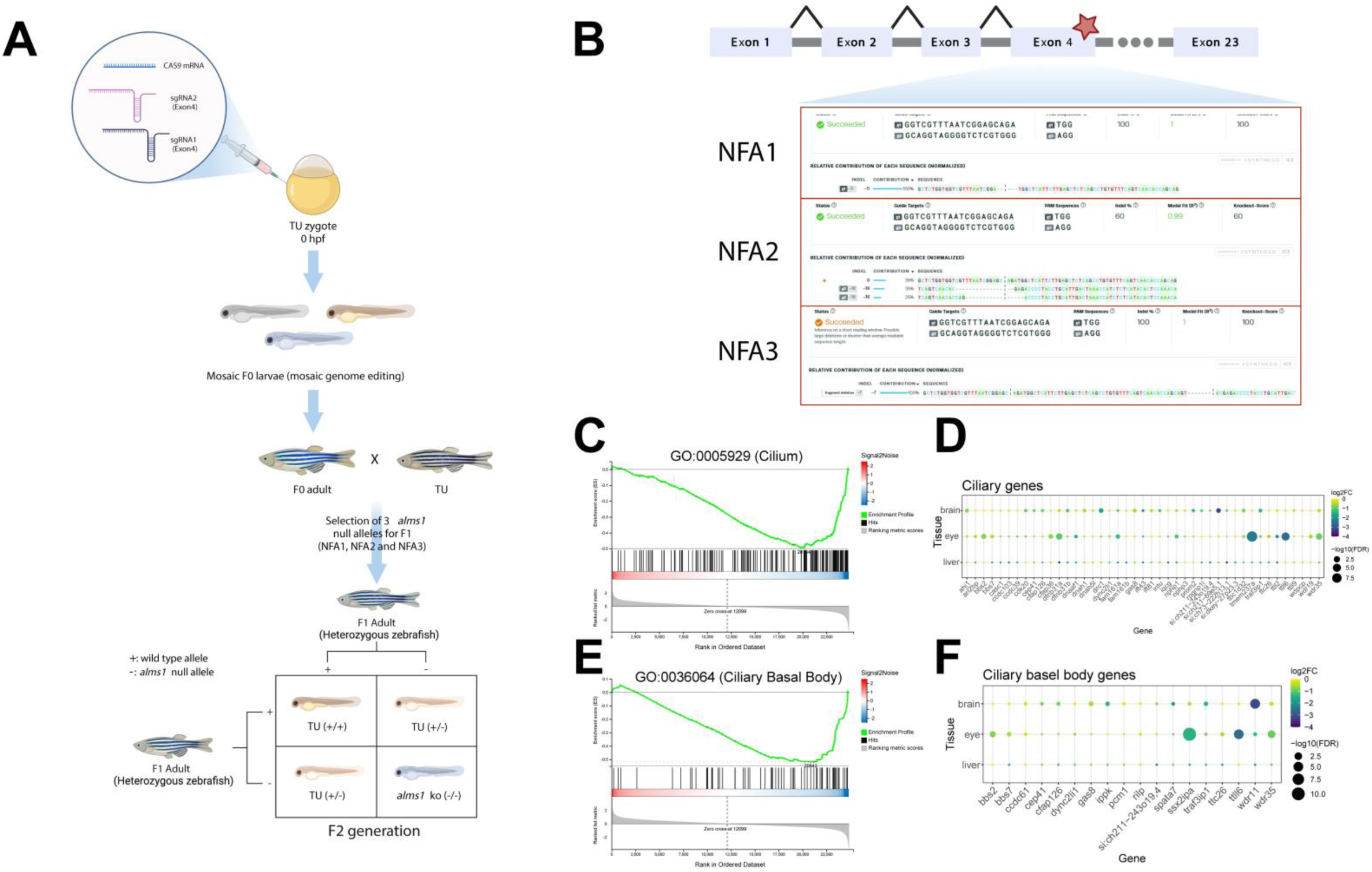
Knockouts generation and validation protocol. **A)** Microinjection and breeding strategy for the zebrafish models generation. **B)** Non-functional alleles (NFAs) of *alms1* characterized after screening. **C)** Gene-set enrichment analysis (GSEA) result for cilia term on zebrafish eyes. **D)** expression levels and significance after differential expression analysis of cilia-associated genes on zebrafish eyes. **E)** Gene-set enrichment analysis (GSEA) result for ciliary basal body term on zebrafish eyes. **F)** Expression levels and significance after differential expression analysis of cilia-basal-body-associated genes on zebrafish eyes.

The theoretically truncated protein would retain the first 9-10% of the full-length protein. In our case, these predicted stop codons do not appear to trigger nonsense-mediated mRNA decay, as suggested by the slight decreases in *alms1* mRNA levels in NFA 1 and 3 **(Figure S3B)**. However, we did observe a marked reduction in ciliary and cilia basal body function in the eyes, the primary ciliated organ analysed in this study **(Figure 1C, E)**. The expression of multiple genes related to these organelles was downregulated **(Figure 1D, F, Figure S4)**. Verifying the reduction in protein levels was not possible, as no zebrafish-specific alms1 antibody is currently available. Previous studies using *alms1* gene knockout models in zebrafish also exhibited these limitations [35].

Moreover, Nesmith et al. reported a reduced number of *alms1 -/-* homozygotes compared to the expected Mendelian ratios, consistent with our findings **(Supplementary Table 4)** [35]. This suggests a high mortality rate in knockout embryos.

### *alms1* depletion has a tissue-dependent impact on zebrafish

The study design included three genotyped wild-type fish (TU) and three *alms1* -/- (2 NFA1 and 1 NFA3) F2 fish **(Figure 2A)**. The quality of the samples was consistent, and, of the total number of genes identified through sequencing, 19,795 were detected in all samples before the application of any expression filters **(Figure 2B)**.

**Figure 2.**
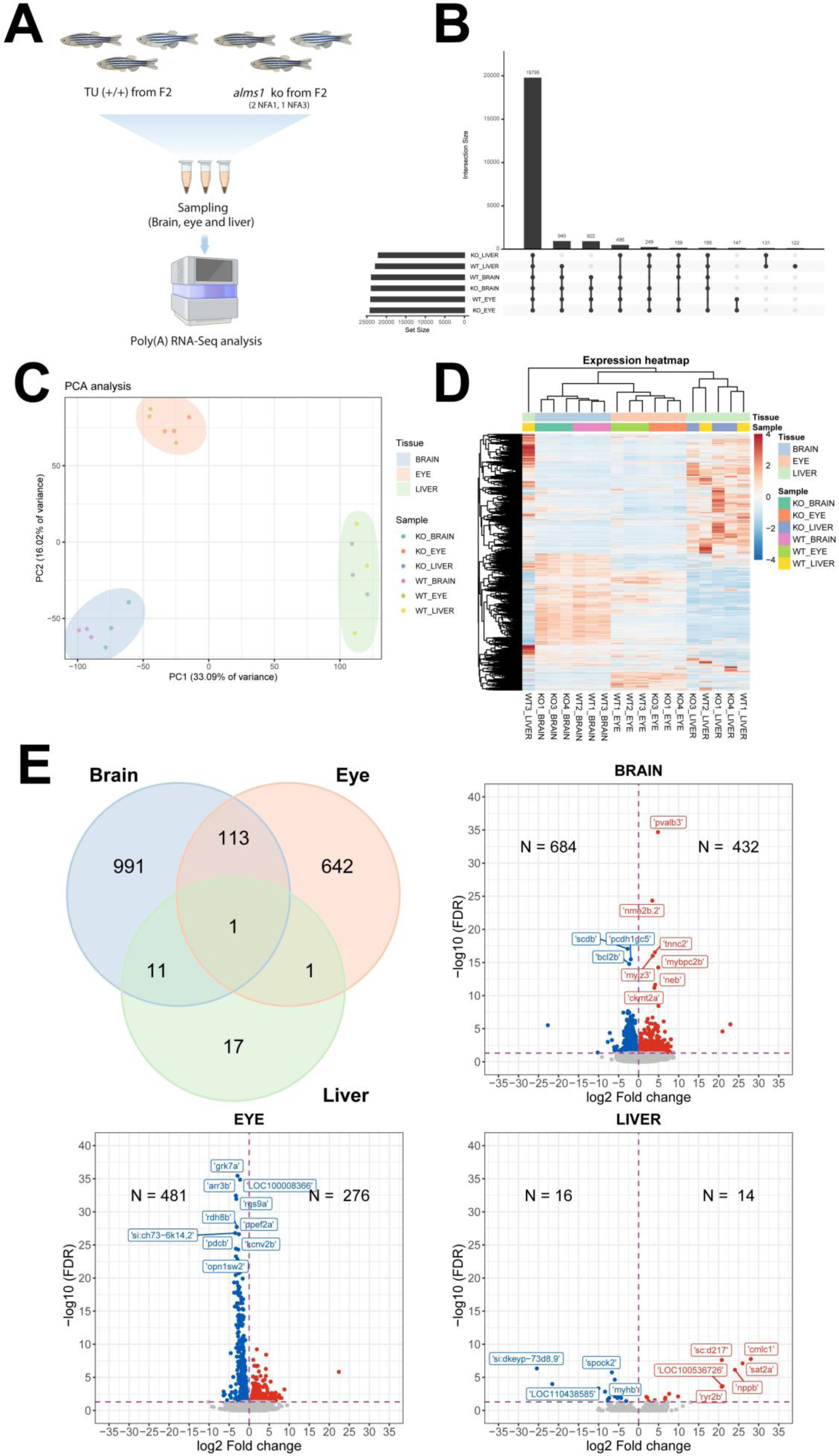
Multi-tissue expression profiles. **A)** Experimental design and protocol for sample collection and sequencing. **B)** Upset plot of the detected genes by sample. **C)** Principal component analysis (PCA) on the various study samples. **D)** Heatmap illustrating the expression profile of each sample. **E)** Tissue overlaps of differentially expressed genes (DEG) and a volcano plot of each tissue, indicating the total number of genes that are overexpressed or underexpressed.

Principal component analysis (PCA) using DESeq2-normalised counts for these common genes revealed that the primary source of variation was the tissue of origin rather than the genotype **(Figure 2C)**. However, within each tissue cluster, samples could be clearly distinguished by genotype in the brain and eye, but not in the liver **(Figure 2D)**. At this stage, only genes with a minimum expression of 30 reads per sample were considered. These genes were subsequently used for downstream analyses.

No phenotypic or expression differences were observed among *alms1-/-* individuals based on the non-functional allele (NFA) they carried **(Figure 2D, Figure S3)**.

Following differential expression analysis, a considerable degree of discrepancy was observed in the extent of overlap between the various tissues. In the case of the brain and eye, the organs where the greatest alterations were detected, the overlap ranges between 10% and 15% **(Figure 2C)**. The number of differentially expressed genes varied substantially across tissues, with the brain exhibiting the most pronounced alterations, comprising 1,116 DEGs (432 overexpressed and 684 underexpressed) **(Figure 2B)**. Notably, only 30 DEGs were identified in the liver, with 14 being overexpressed and 16 being underexpressed, despite the liver being the organ from which the highest amount of RNA was extracted **(Figure 2C)**. This highlights the tissue-specific regulatory function of the *alms1* gene **(Figure 2C)**.

### Glutamatergic synapses as a convergent alteration between the eyes and the brain in response to *alms1* depletion

A preliminary over-representation analysis (ORA) using the Kyoto Encyclopaedia of Genes and Genomes (KEGG) pathways revealed the existence of both tissue-specific and common processes **(Figure 3)**. In the eyes, the most representative process was identified as phototransduction. In the brain, processes related to infection and, consequently, alterations of the innate immune system, as well as synaptic processes, were highlighted **(Figure 3A, B, Figure S5)**. In the liver, only a few significant processes were identified, and these did not appear to follow any discernible pattern. This may be attributed to the limited number of alterations detected in the differential expression analysis. **(Figure 3C)**. Consequently, no further analysis was carried out with the results of this organ.

**Figure 3.**
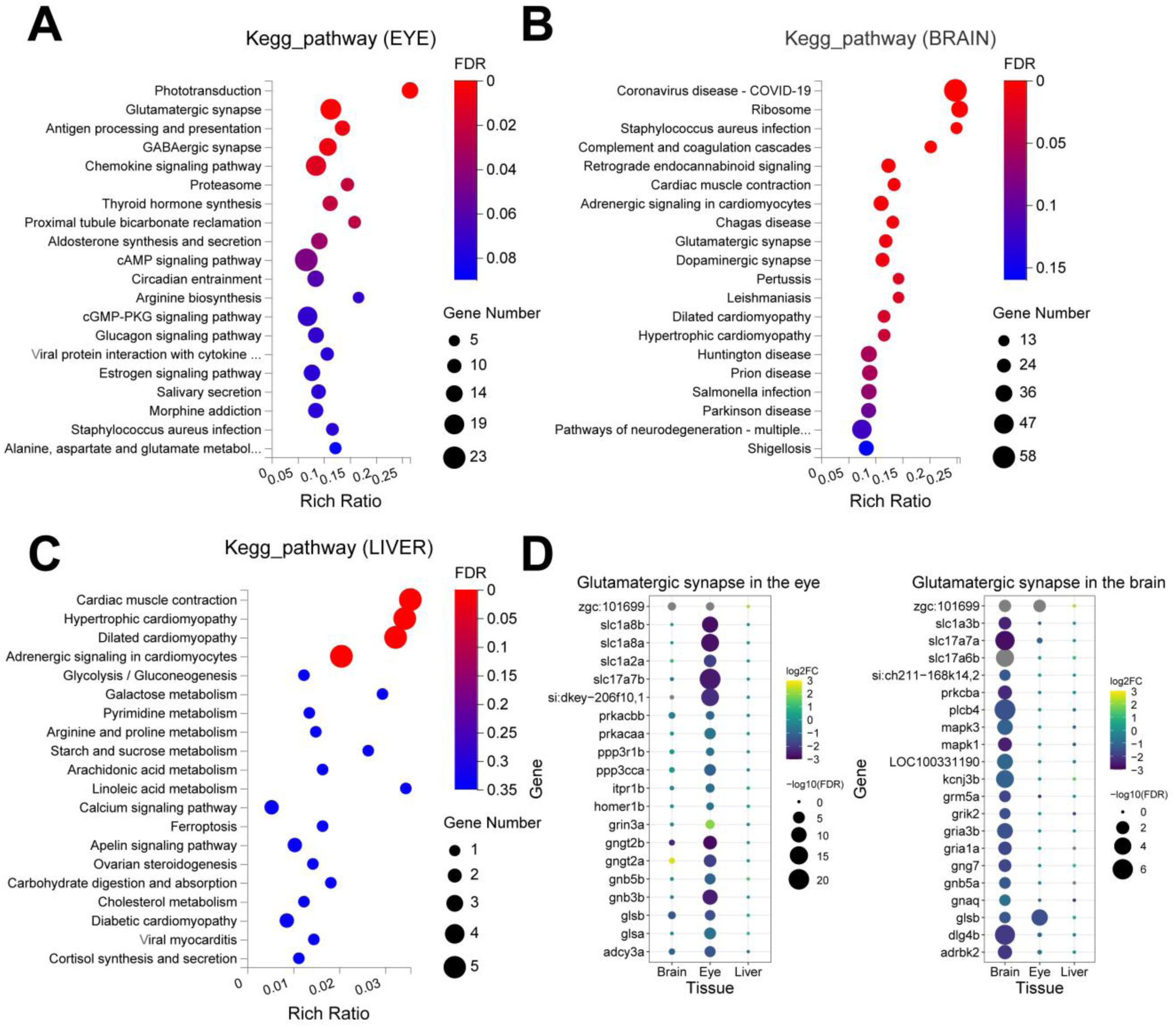
Multi-tissue over-representation analysis (ORA). **A)** ORA of KEGG pathways in differentially expressed genes in the eye. **B)** ORA of KEGG pathways in differentially expressed genes in the brain. **C)** ORA of KEGG pathways in differentially expressed genes in the eye. **D)** Expression profiles of glutamatergic synapse-associated genes in the eye and brain.

A noteworthy process common to both the brain and eye results is the glutamatergic synapse (**Figure 3A, B)**. Although this process is significantly represented in both organs, the associated differentially expressed genes (DEGs) did not overlap **(Figure 3D)**. This underscores the tissue-specific regulation of the *alms1* gene, even within the same biological process.

### Inflammation and the innate immune alterations serve as the connecting link between eye and brain disorders

The results of the expression analysis indicated that *alms1 -/-* individuals exhibited an elevated chemotactic response of neutrophils and other leukocyte, particularly B lymphocytes, alongside a pronounced reaction to type II interferon **(Figure 4A)**. Conversely, the inhibited processes were primarily associated with ciliary, sensory, and nervous functions specific to the eye and its photoreceptors **(Figure 4A)**. The sensory dysfunction of the eye and its photoreceptors appears to be primarily linked to genes belonging to the opsin family and G proteins **(Figure 4B)**.

**Figure 4.**
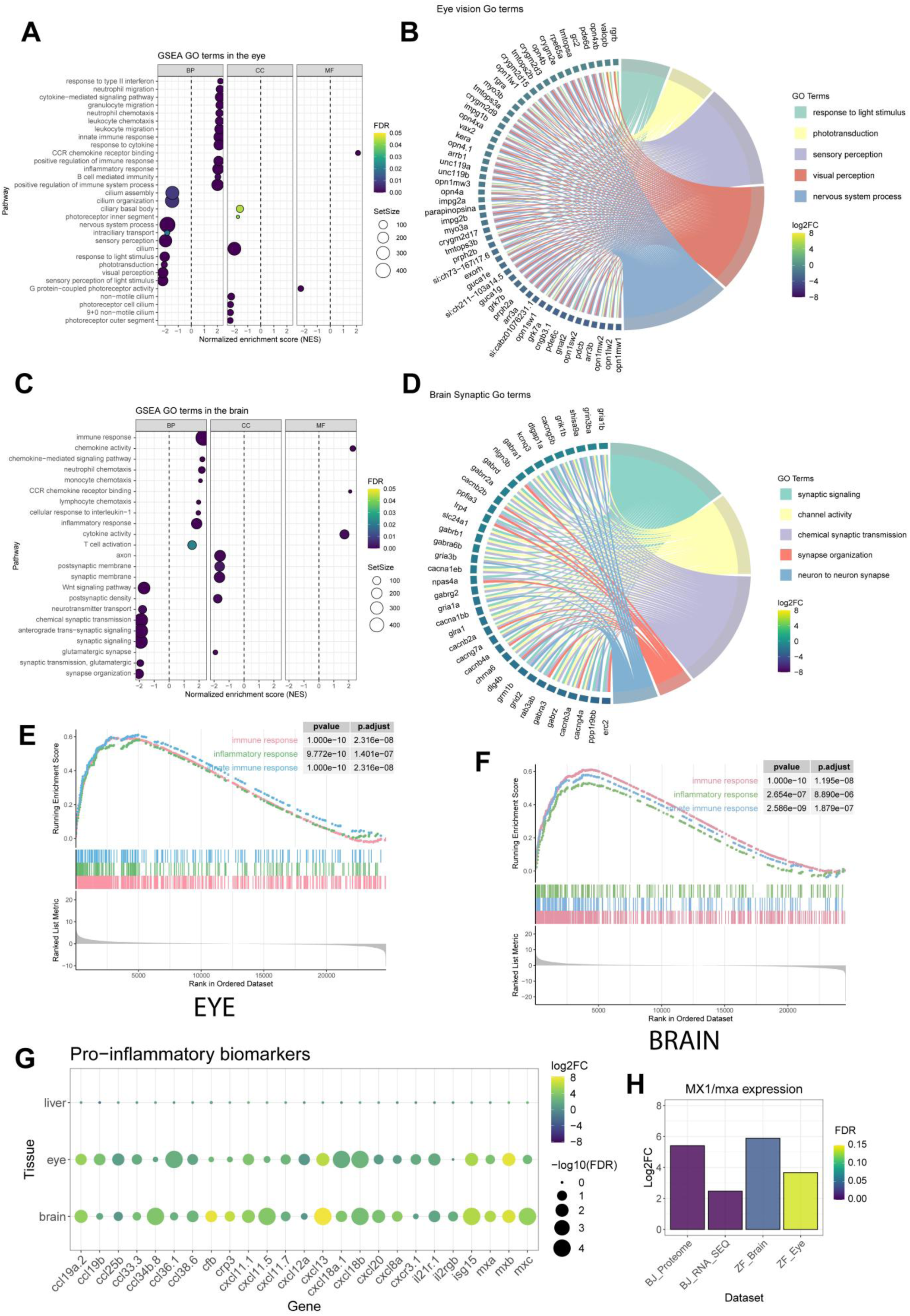
Multi-tissue gene set enrichment analysis (GSEA). **A)** Normalised enrichment scores of different eye GO terms. **B)** Chord plot of genes related to the main processes involved in ocular dysfunction. **C)** Normalised enrichment scores of different brain GO terms. **D)** Chord plot of genes related to the main processes involved in brain dysfunction. **E)** GSEA-plot of immune response in the eye. **F)** GSEA-plot of immune response in the brain. **G)** Expression profile of various pro-inflammatory biomarkers in the different tissues of the study. **H)** Levels of *MX1*/*mxa* gene overexpression in human cell models and zebrafish model.

Conversely, the brain expression profile demonstrated an immune response predominantly mediated by neutrophils, monocytes, and T lymphocytes **(Figure 4C)**. In this organ, functional inhibition was primarily associated with a reduction of synaptic processes mediated by neurotransmitter transport **(Figure 4C)**. This dysfunction in synaptic processes appears to be mainly linked to a decreased expression of GABA receptors, potassium (K^+^) channels and calcium (Ca^2+^) channels **(Figure 4D)**.

Inflammation and overactivation of the innate immune response are common occurrences in both the eye and brain of our models **(Figure 4 E, F)**. In this context, the overexpression of various inflammatory biomarkers was consistently observed across different tissues **(Figure 4G)**. Furthermore, the overexpression of multiple cytokines of the *ccl* and *cxcl* families was observed, along with interleukin-2 and −21 receptors (*il21r* and *il2r*) **(Figure 4G)**. Additionally, the overexpression of *isg15* and various paralogues of the *mx* family in zebrafish (*mxa*, *mxb* and *mxc*), which are markers of interferon pathway activation, is particularly noteworthy **(Figure 4G)**. Notably, the human orthologue of the *mxa* gene is *MX1*, which has already been shown to exhibit marked overexpression in our human cell model in h-TERT fibroblasts **(Figure 4H)** [33].

### Depletion of ciliary genes induces expression signatures associated with accelerated brain ageing

We selected genes related to cognitive processes to further investigate the brain alterations observed in our zebrafish model and determine whether these could be linked to similar alterations in humans. Then, we mapped them to their corresponding human orthologues. Subsequently, an enrichment analysis was performed using the DISGENET disease database.

At the disease level, the human orthologues were found to be associated with a range of brain pathologies, including schizophrenia, bipolar disorder, and intellectual disability **(Figure 5A)**. At the process level, we observed significant enrichment in neurogenesis, neurodevelopment, glutamatergic and dopaminergic synapses and neuroinflammation **(Figure 5B)**.

**Figure 5.**
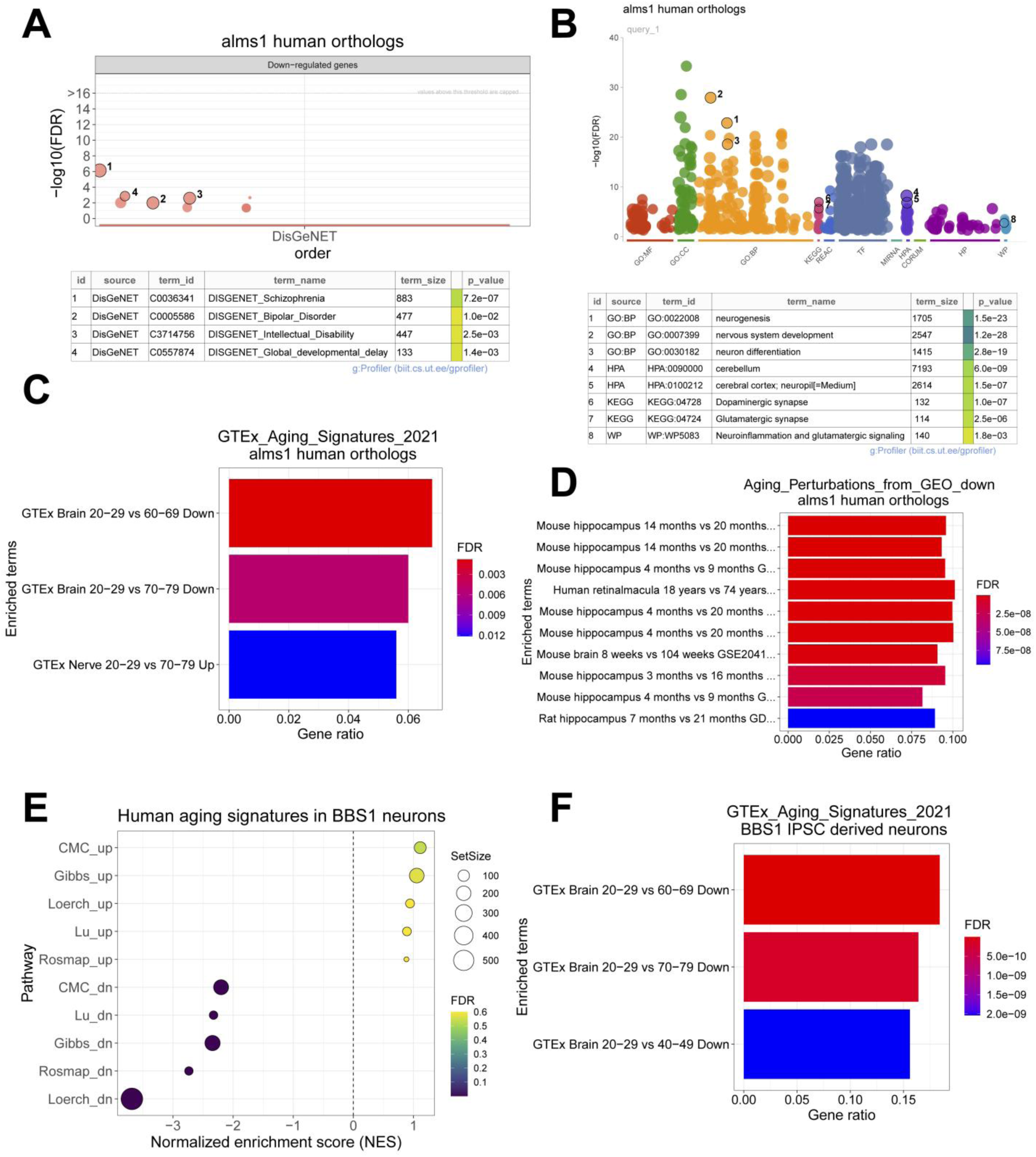
A detailed examination of the expression signatures in the brain. **A)** ORA in DISGENET of *ALMS1* human orthologues. **B)** ORA for GO terms of *ALMS1* human orthologues. **C)** GTEX enrichment analysis for the *ALMS1* gene. **D)** GEO mouse ageing perturbation enrichment analysis for the *ALMS1* gene. **E)** Enrichment of human ageing signatures in *BBS1* neurons. **F)** GTEX enrichment analysis for the *BBS1* gene.

Upon testing the significance of the gene expression signature composed of human brain orthologues, with ageing signatures inferred from the GTEX database, we observed a significant enrichment in genes that are normally differentially expressed when comparing brains from adults aged 20-29 years with those from adults aged 60-69 or 70-79 years **(Figure 5C)**. The same results were observed for the mouse hippocampus **(Figure 5D)**.

To determine whether the enrichment in ageing-associated expression signatures was exclusive to the *ALMS1* gene or could also occur in other ciliary genes, we conducted similar analyses on a *Bardet-Biedl syndrome 1* (*BBS1*) *gene* model. The dataset was generated from neurons derived from induced pluripotent stem cells (iPSCs) of patients with Bardet-Biedl syndrome [45]. The patients in question carried pathogenic variants in the *BBS1* gene.

The DEGs from the neurons of these patients were used in an enrichment analysis with gene expression signatures obtained from previously published cohorts. These cohorts have been used to assess brain ageing and the onset of neurodegenerative diseases and were previously curated and published in Mavrikaki *et al* 2022 **(Figure 5E)** [46]. The analysis yielded significant results for all inhibitory expression signatures across all cohorts **(Figure 5E)**.

Finally, DEGs from *BBS1* neurons were analysed against the GTEX-inferred ageing dataset, yielding results similar to those observed in the *ALMS1* dataset but with a higher level of significance **(Figure 5F)**.

## DISCUSSION

In this study, multi-tissue expression profiling was conducted in zebrafish models with *alms1* gene depletion. Exon 4 was selected as the target for gene editing in our model, as previous studies have demonstrated its role in recapitulating the Alström syndrome (ALMS) phenotype in zebrafish [35]. The high mortality observed in our *alms1* knockout homozygous embryos was consistent with that described in the literature **(Supplementary Table 4)** [35].

During the optimization of the technique, several tissues associated with the ALMS phenotype were initially identified as potential candidates for RNA extraction. However, the study ultimately focused on the eye, brain, and liver. These tissues were deemed the most relevant for elucidating the underlying pathology, as they provided sufficient RNA quality for sequencing. For future studies, we recommend incorporating pooled samples from additional organs not included in this selection, such as the heart, pancreas, and gut.

The most prevalent phenotype associated with ALMS is visual impairment [3,47]. Previous studies with *alms1* morpholinos in zebrafish larvae had already pointed out phototransduction, detection of light stimulus, and visual perception as some of the most altered processes when inhibiting the *alms1* gene [48]. Our results confirm that these phenotypes are not transient but remain stable following the depletion of the *alms1* gene in zebrafish **(Figure 2E, 3A and 4A)**. This aligns with a more recent study that confirmed, through histological analysis, that *alms1* gene knockout models in zebrafish exhibit retinal dystrophy [44].

On the other hand, brain involvement is not uncommon in ALMS [49,50]. it is plausible that both metabolic syndrome and hearing loss are linked to brain disorders and other less common phenotypes, such as mental disorders [3,47]. This theory is further supported by our data, which reveal the major alterations in the brain **(Figure 2E),** along with other studies that have reported no changes in hair cells [37]. In addition, it has also been described that deficits in auditory working memory are common in patients with ALMS [50]. Thus, the hearing loss presented by these patients seems to be due more to a brain dysfunction than to a problem in the ears themselves, although they have hair cells. Previous studies in zebrafish larvae reported neuronal alterations upon *bbs1* inhibition but not *alms1* [48]. In light of new data from our study, this suggests that neuronal alterations in ALMS may emerge later than in Bardet-Biedl Syndrome (BBS), another ciliopathy closely related to ALMS.

Liver fibrosis is also a relatively common phenotype in ALMS, yet this organ has been little studied to date in ALMS animal models [3,27]. In this context, given that the zebrafish has been validated as a model organism for studying human liver diseases, our findings suggest that the observed liver alterations are a secondary symptom, rather than a direct consequence of the gene’s absence in the tissue **(Figure 2E)** [51,52].

The initial finding relates to the apparent tissue-dependent regulation of this gene. The hypothesis of tissue-dependent regulation in ALMS and the existence of isoforms with distinct functions has long been a subject of interest in the study of this pathology [3]. Nevertheless, to date, in vitro or animal model studies on this syndrome have only focused on single cell type, tissue, or organ. To our knowledge, this is the first study to integrate the gene expression signature across multiple whole organs in an animal model of ALMS, confirming the pleiotropic role of the *ALMS1* gene and thus providing a valuable resource for future research.

As previously discussed, the most pronounced alterations were observed in the brain among the different tissues **(Figure 2D)**, which also exhibited the highest levels of *alms1* expression across the various organs **(Figure S3B)**. Furthermore, a significant correlation was observed between the relative expression of *alms1* and the extent of tissue alterations **(Figure 2D; Figure S3B)**. For instance, in the liver, minimal *alms1* expression was detected and no notable alterations in the expression profile were observed.

In both the eye and the brain, the processes identified as inhibited in the enrichment analyses were associated with the dysfunction of each organ, respectively **(Figure 4A-B, C-D)**. However, the overexpressed processes appear to share a common origin, namely the interferon-mediated inflammatory response primarily driven by the innate immune system **(Figure 4)**.

These alterations appear to result from dysregulated self-immunity and self-tolerance. This phenomenon has previously been observed BBS [19]. Similar alterations are relatively common in another group of rare diseases known as type I interferonopathies. This group comprises mainly monogenic autoinflammatory mendelian disorders, primarily associated with mutations in TREX1, RNASEH2A, RNASEH2B, RNASEH2C, SAMHD1, ADAR, IFIH1, TMEM173, ACP5, ISG15 and DDX58, as well as early components of the complement cascade, particularly C1 and C4 [53].

Type I IFNs are typically induced by viral pathogens. Viral infections frequently involve host-encoded nucleic acid-binding pattern-recognition receptors [53]. This is likely the primary reason why over-represented pathways in the brain were associated with infectious diseases **(Figure 3B, Figure S5)**. One of the most prevalent symptoms observed in Type I interferonopathies is neuroinflammation, which appears to be present in the brains of our zebrafish model and could potentially occur in humans as well **(Figure 4F, 5B)** [54]. In light of these observations, it seems reasonable to hypothesise that several of the common neurological alterations in ALMS may be attributable to, or influenced by, chronic brain inflammation [3]. Furthermore, chronic neuroinflammation has been linked to other neurodegenerative diseases, including Alzheimer’s disease and Schizophrenia. Certain human orthologues identified in our model appear to be associated with the latter [55,56]. It is plausible that this progressive brain dysfunction, driven by chronic inflammation, may underlie the emergence of gene expression signatures associated with accelerated brain ageing in models of ciliopathies such as ALMS or BBS **(Figure 5C-F)**.

This brain dysfunction may also be a significant contributing factor to, or a consequence of, the metabolic syndrome that characterises these patients [3]. Determining which of the two processes occurs first is challenging, as both neurological (hearing and vision loss) and metabolic (obesity and type 2 diabetes mellitus) disorders manifest at an early age in the majority of cases and persist throughout the patient’s lifetime [23,47]. The *GLUT4* gene, which is associated with alterations in adipose tissue glucose transport and insulin resistance in Alström syndrome, also plays a critical role in the brain. This is evidenced by its function in ensuring the brain’s glucose requirements and its involvement in cognitive processes such as memory [26,28,29].

Several studies have reported the loss of β-cells in the pancreas following the deletion of the *alms1* gene in zebrafish [35,38]. This leads to impaired glucose sensitivity, and it is plausible that a similar deficiency also occurs in the brain. This occurs in the context of overexpression of the transcription factor *NEUROD1*, an apoptosis protector in pancreatic β-cells, which appears to be a compensatory mechanism [35]. This presumed lack of glucose sensitivity could lead to neuroglycopenia, a state of cerebral hypoglycaemia characterised by increased oxidative stress and the induction of chronic inflammation mediated by the innate immune system [57–59]. Thus, the neuroinflammation observed in our model could be a consequence of ALMS metabolic syndrome. This neuroinflammation, along with the oxidative stress induced by hypoglycaemia, may contribute to chronic stress in the brain, leading to a progressive decline in cognitive functions at a faster rate than expected in a healthy individual **(Figure 5)** [59].

It has previously been postulated that ciliopathies may serve as a model disease for accelerated ageing pathology. In particular, our previous meta-analysis of 227 patients described in the literature revealed a correlation between syndromic score and age, with an accelerated progression during the first two decades of life [47]. Subsequently, this progression appeared to stabilise.

It seems reasonable to assume that the *ALMS1* gene, which regulates cell migration, division and apoptosis, may also play a role in ageing [9,33,60,61]. However, the precise mechanisms underlying this process remain unclear, and further research is required to elucidate the role of *ALMS1*, the centrosome, and the primary cilium in ageing.

## CONCLUSIONS

This study is the first to demonstrate the role of the *alms1* gene in regulating gene expression across multiple tissues, highlighting its tissue-specific regulatory functions. Furthermore, a link between *alms1* gene depletion and chronic ocular and brain dysfunction was established. The most significant transcriptomic alterations were observed in the brain and eyes, characterised by ciliary dysfunction and a dysregulated immune response involving innate and adaptive immunity, including the activation of T and B lymphocytes. Chronic inflammation, particularly neuroinflammation, was associated with disrupted synaptic processes, including glutamatergic and dopaminergic synapses, and reduced expression of GABA receptors and ion channels. These findings link *alms1* depletion, cognitive dysfunction, and neuro-ocular inflammation.

The observed dysfunction appears to give rise to a notable age-related expression signature in the brain. Furthermore, this ageing-related expression signature also seems to be present in previously published datasets for other ciliopathies, such as Bardet-Biedl syndrome, where autoimmune alterations have also been reported. This suggests that, in addition to the *ALMS1* gene, the primary cilium and centrosome are organelles that potentially regulate immunity and ageing.

## Author Contributions

BB-M, DV, JR and PS-B designed the study. BB-M, LM-M, CC and LG-P performed the experiments. BB-M performed data analysis and interpretation of results. BB-M wrote the draft manuscript. All authors reviewed the manuscript and provided approval for publication.

## Informed Consent Statement

Not Apply.

## Data Availability Statement

Data are available via Sequence Read Archive (SRA) with the identifier PRJNA1208400

## Code Availability Statement

The code used in this study can be accessed at the GitHub address https://github.com/bmascat/rna-seq-alms-zebrafish.

## Acknowledgements

We sincerely thank the Genomics services from Centro de Apoyo Científico-Tecnológico a la Investigación (CACTI) of the University of Vigo and its specialists Ángel Sebastián Comesaña, Verónica Outeiriño for them for its work on sequencing of genetic screening samples. We also thank the Spanish Association of Alström syndrome patients for its technical and financial support

