## Supplementary material for "*alms1* regulates the immune response and brain ageing in zebrafish"

CINBIO Facultad de Biología, Universidad de Vigo, Campus As Lagoas-Marcosende s/n, 36310 Vigo, Spain

### Supplementary Tables:

**Supplementary Table 1.** Sequences of microinjected gRNAs and primers used to amplify exon 4 of the *alms1* gene in zebrafish.

| ID | Sequence |
|---|---|
| gRNA 1 | aattaatacgactcactata**GGTCGTTTAATCGGAGCAGA**gttttagagctagaaatagc |
| gRNA 2 | aattaatacgactcactata**GCAGGTAGGGGTCTCGTGGG**gttttagagctagaaatagc |
| CRISPR-Scaffold | gatccgcaccgactcggtgccactttttcaagttgataacggactagccttattttaacttgctatttctagctctaaaac |
| alms1-Exon4-FWD | TGGATACACACCAAACAGCAGA |
| alms1-Exon4-RVS | TCTGGACTGGTTGCTTTGAGG |

**Supplementary Table 2.** Sensitivity, specificity and localisation parameters of the gRNAs of the study. The reference transcript used was *alms1* (ENSDARG00000074779).

| ID | Position | Strand | Exon | gRNA sequence | PAM | Specificity | Efficiency | Reference |
|---|---|---|---|---|---|---|---|---|
| gRNA 1 | 9041962 | + | 4 | GGTCGTTTAATCGGAGCAGA | TGG | 93 | 59 | this study |
| gRNA 2 | 9042020 | - | 4 | GCAGGTAGGGGTCTCGTGGG | AGG | 96 | 62 | this study |

**Supplementary Table 3.** Non-functional alleles (NFA) selected in F1 to establish homozygous F2. Length of the insertion-deletion (Indel) that was generated, type of mutation, knock-out score (KO), calculated by the ICE programme and gRNA was responsible for the deletion.

| ID | Sequence | Indel (pb) | Type | KO score | gRNA |
|---|---|---|---|---|---|
| NFA1 | GCTCTGGTGGTCGTTTAATCGGA--\|---TGGCTCATTCTTGAGCTCTCAGC | -5 | Frameshift | 100 | 1 |
| NFA2 | GCCTGTGTTTCAGTCAACAC--------------\|--GAGACCCCTACCTGCATTG | -16 | Frameshift | 100 | 2 |
| NFA3 | GCCTGTGTTTCAGTCAACACCAGCAGT-------\|ACGAGACCCCTACCTGC | -7 | Frameshift | 100 | 2 |

**Supplementary Table 4.** Absolute frequencies of each genotype per non-functional allele characterised at F2. *: one of the NFA3 homozygous individuals died before it could be sacrificed for dissection.

| **F2 Screening** | NFA1 | NFA2 | NFA3 |
|---|---|---|---|
| Homozygous | 3 | 0 | 2 |
| Heterozygous | 23 | 10 | 18 |
| Wild-type | 9 | 4 | 3 |
| **Total** | **35** | **14** | **23** |

### Supplementary Figures:

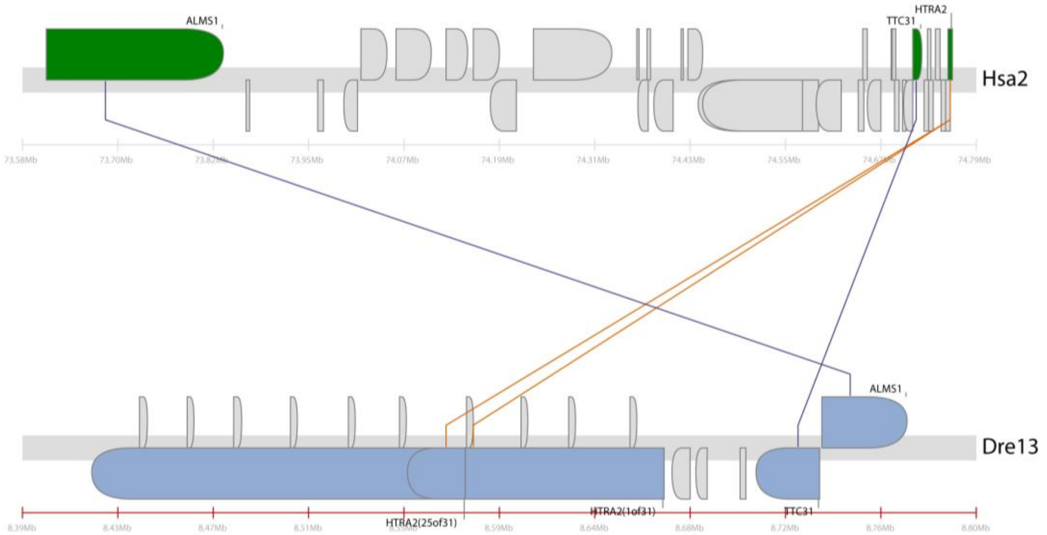


**Supplementary Figure S1.** Parallelism and structural comparison between the *ALMS1* gene in humans, chromosome 2 (Hsa2) and zebrafish, chromosome 13 (Dre13).

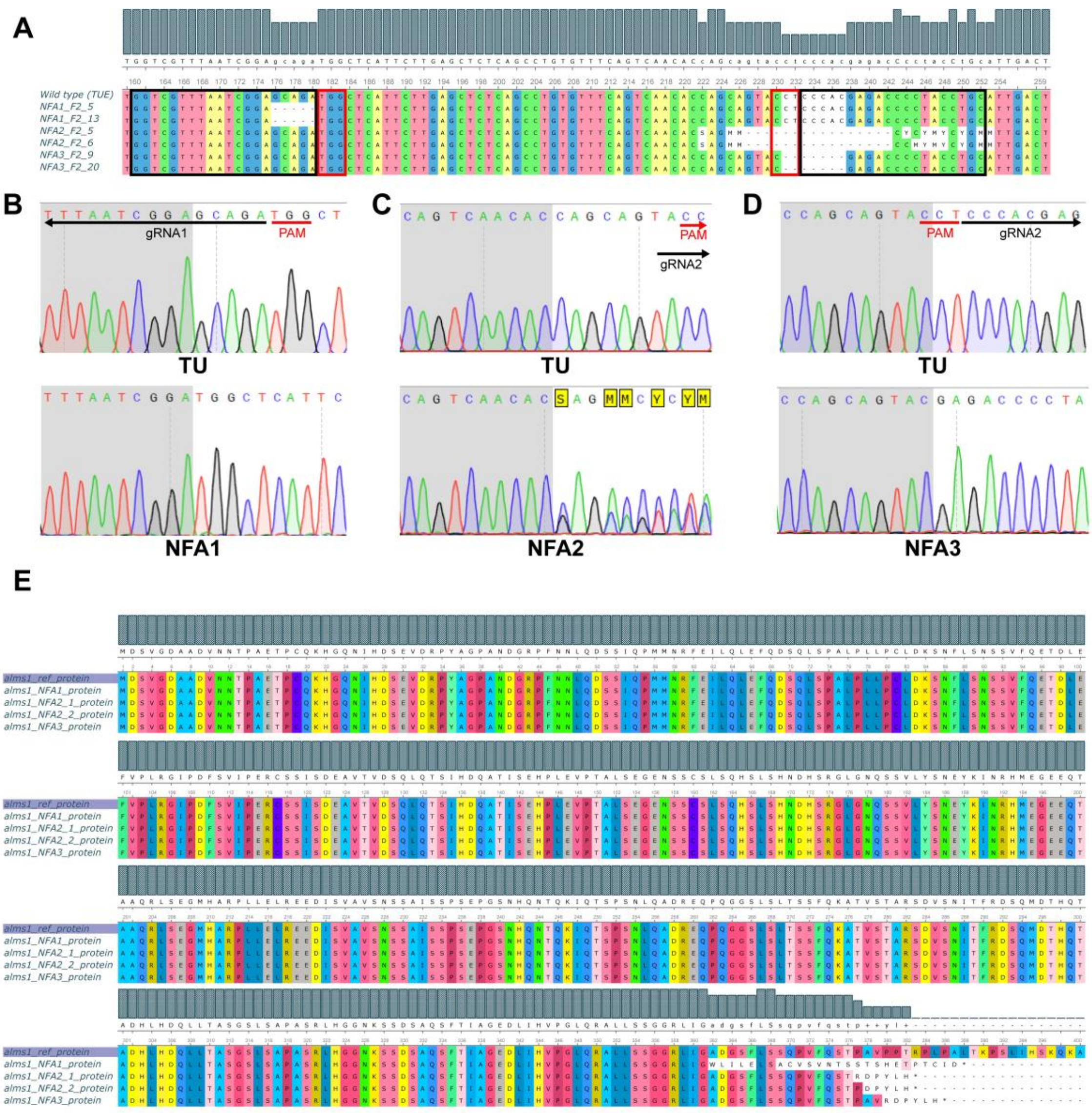


**Supplementary Figure S2.** Genotyping of study individuals. **A)** Multi-sequence alignment (MSA) of cDNA from exon 4 of individuals carrying each genotype. **B), C), D)** sequence cut-off point for each of the non-functional alleles (NFA) compared to the wild-type genotype. **E)** Prediction of amino acid sequence resulting from mutation of each allele

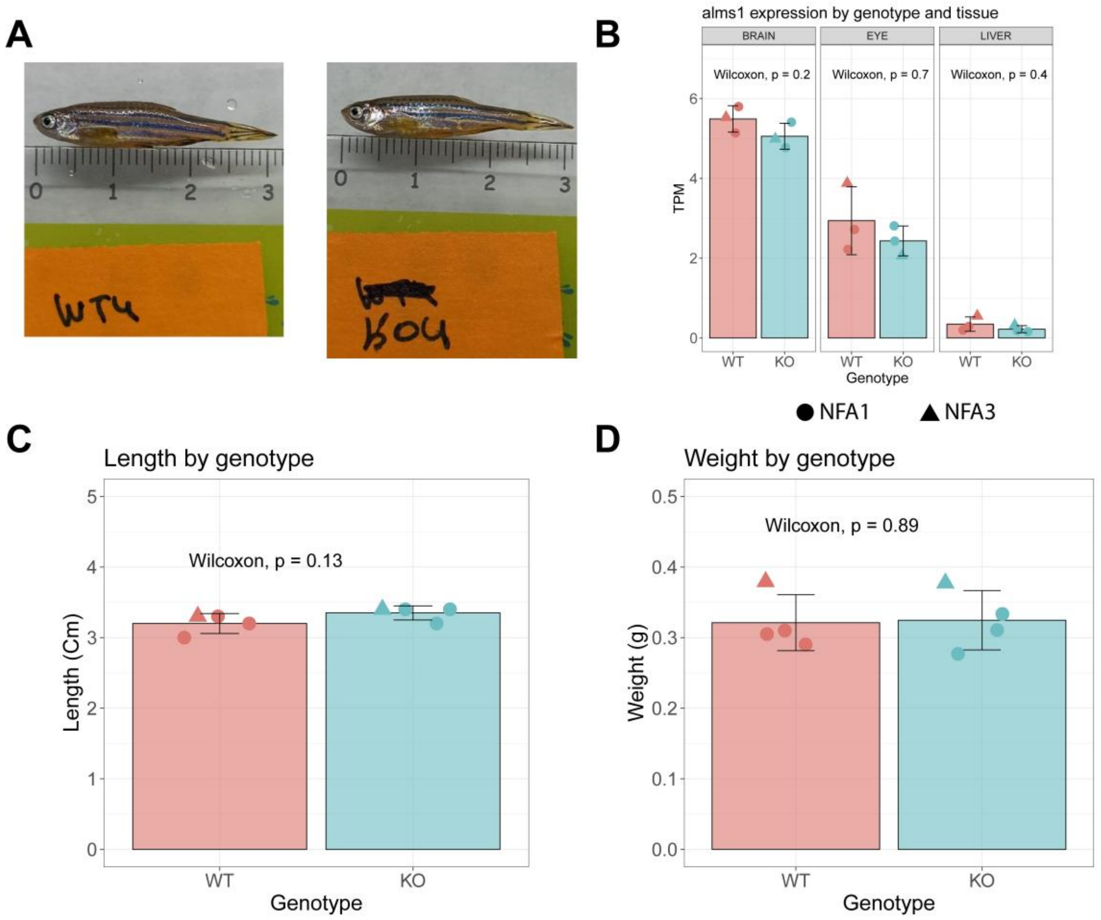


**Supplementary Figure S3.** Comparison of wild type (TU) vs. *alms1* ko phenotypes. **A)** Visual inspection of individuals with different genotypes. **B)** Level of RNA expression normalised as transcripts per million (TPM) for the *alms1* gene in the different tissues of the study. **C)** Comparison of lengths between individuals. **D)** Comparison of weights between individuals.

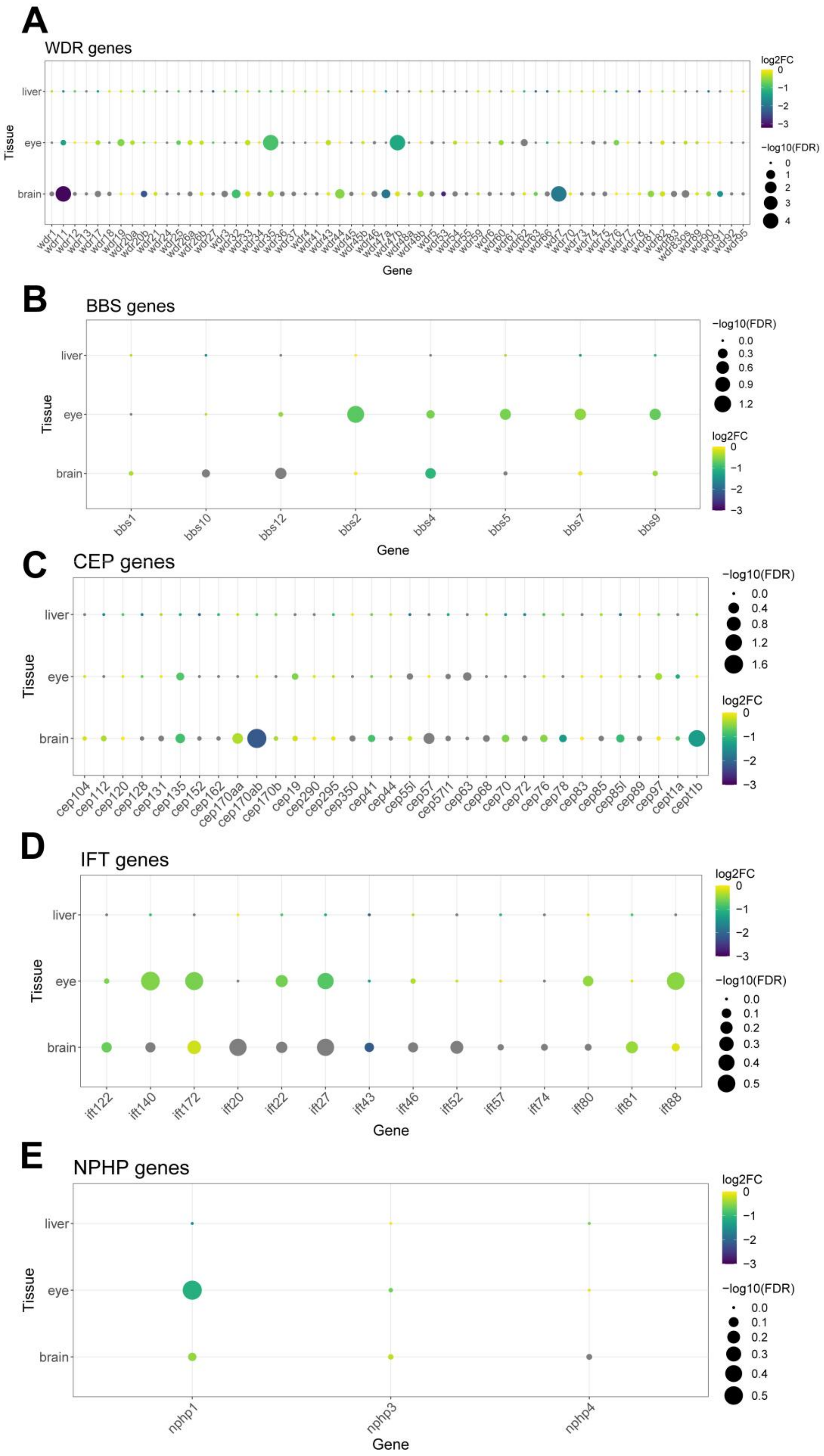
A
WDR genes
log2FC
0
−1
−2
−3
−log10(FDR)
0
1
2
3
4
liver
eye
brain
Tissue
wdr1
wdr11
wdr12
wdr13
wdr17
wdr18
wdr19
wdr20a
wdr20b
wdr21
wdr24
wdr25
wdr26a
wdr26b
wdr27
wdr3
wdr32
wdr33
wdr34
wdr35
wdr36
wdr37
wdr4
wdr41
wdr43
wdr44
wdr45
wdr45b
wdr46
wdr47a
wdr47b
wdr48a
wdr48b
wdr5
wdr53
wdr54
wdr55
wdr59
wdr6
wdr60
wdr61
wdr62
wdr63
wdr66
wdr7
wdr70
wdr73
wdr74
wdr75
wdr76
wdr77
wdr78
wdr81
wdr82
wdr83
wdr83os
wdr89
wdr90
wdr91
wdr92
wdr95
Gene
B
BBS genes
−log10(FDR)
0.0
0.3
0.6
0.9
1.2
log2FC
0
−1
−2
−3
liver
eye
brain
Tissue
bbs1
bbs10
bbs12
bbs2
bbs4
bbs5
bbs7
bbs9
Gene
C
CEP genes
−log10(FDR)
0.0
0.4
0.8
1.2
1.6
log2FC
0
−1
−2
−3
liver
eye
brain
Tissue
cep104
cep112
cep120
cep128
cep131
cep135
cep152
cep162
cep170aa
cep170ab
cep170b
cep19
cep290
cep295
cep350
cep41
cep44
cep55l
cep57
cep57l1
cep63
cep68
cep70
cep72
cep76
cep78
cep83
cep85
cep85l
cep89
cep97
cept1a
cept1b
D
IFT genes
log2FC
0
−1
−2
−3
−log10(FDR)
0.0
0.1
0.2
0.3
0.4
0.5
liver
eye
brain
Tissue
ift122
ift140
ift172
ift20
ift22
ift27
ift43
ift46
ift52
ift57
ift74
ift80
ift81
ift88
Gene
E
NPHP genes
log2FC
0
−1
−2
−3
−log10(FDR)
0.0
0.1
0.2
0.3
0.4
0.5
liver
eye
brain
Tissue
nphp1
nphp3
nphp4
Gene

**Supplementary Figure S4.** Expression levels in our study individuals of different gene families related to the primary cilium and ciliopathies. **A)** WD repeat domain (WRD) family genes. **B)** Bardet Biedl syndrome (BBS) family genes. **C)** Centrosome and Cytoskeleton Protein (CEP) family genes. **D)** Intra-flagellar transport family (IFT) genes. **E)** Nephronophthisis family genes (NPHP).

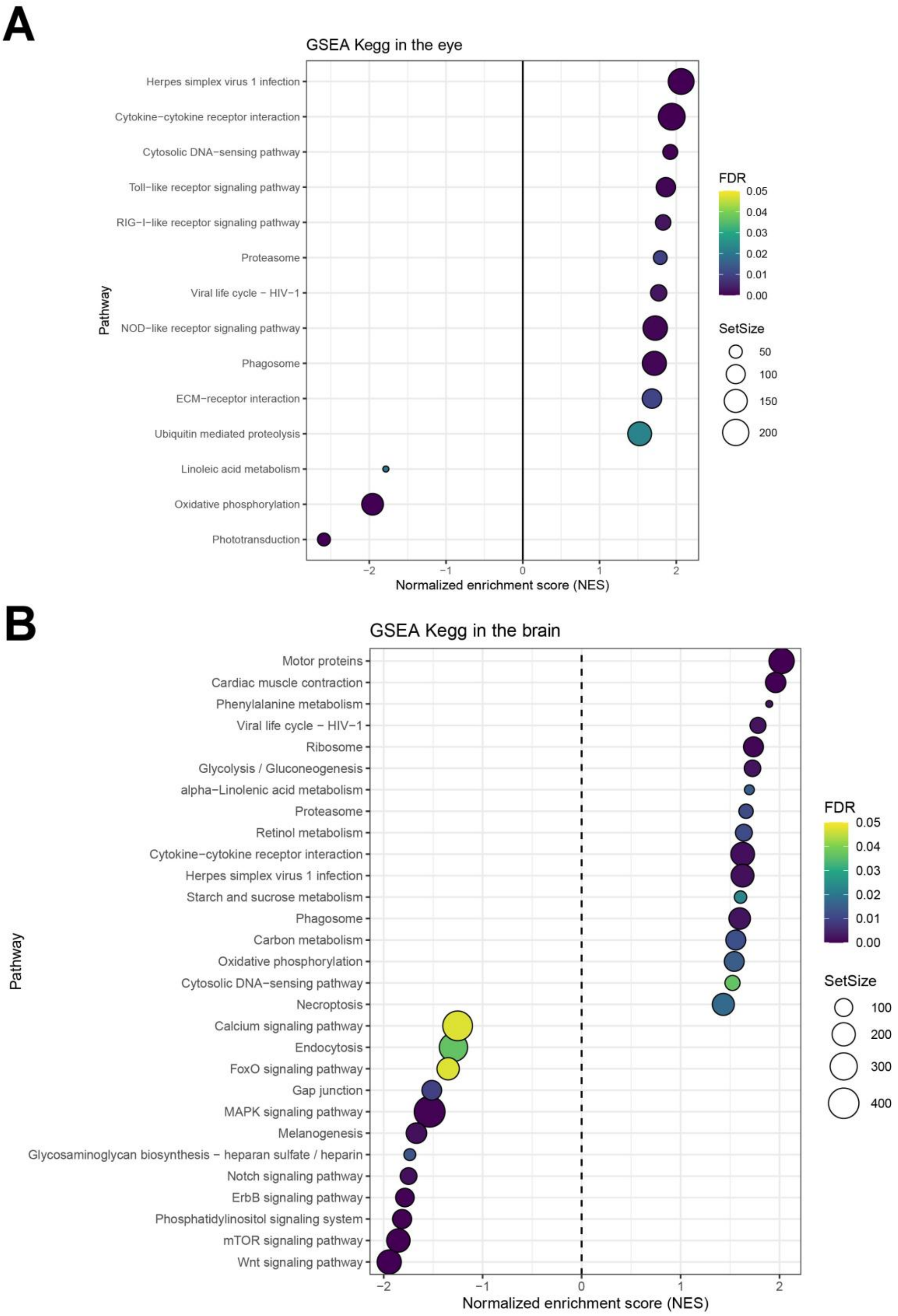


**Supplementary Figure S5.** Gene-Set Enrichment Analysis (GSEA) in Kegg pathways. **A)** Normalised enrichment score (NES) on statistically significant processes in eye. **B)** Normalised enrichment score (NES) on statistically significant processes in brain.
